# Contact-dependent growth inhibition drives kin discrimination and shapes in vivo strain dominance in *Pseudomonas aeruginosa* infection

**DOI:** 10.64898/2026.09.30.755834

**Authors:** Natasha Peterson, Brandi L. Cobe, Adam S. Bronson, Jonathan P. Allen

## Abstract

*Pseudomonas aeruginosa* is a ubiquitous environmental bacterium and an opportunistic pathogen. Rather than existing as single clonal lineages, environmental and clinical populations often consist of a mixture of distinct strains that directly compete to dominate the niche. To eliminate rivals, *P. aeruginosa* deploys a form of interference competition termed contact-dependent growth inhibition (CDI). CDI is mediated by CdiA, where, upon cell contact, the CdiA C-terminal toxin domain is delivered to prey cells. Toxin-producing cells encode for a cognate CdiI immunity protein to prevent self-intoxication. Different strains of *P. aeruginosa* encode for CdiA proteins with different C-terminal toxin domains and cognate CdiI immunity proteins. Most studies have relied on engineered strains to investigate the mechanism of CDI. While extremely useful for studying CdiA-mediated intoxication, such work masks the true diversity of wild-type CdiA variants and their impact on competitive interactions between strains. In this study, we systematically investigated CDI-mediated competition using unique conspecific strains of *P. aeruginosa*. We identified several novel CdiA toxin domains and mapped the competitive outcomes they govern. Furthermore, using a mouse model of acute pneumonia, we demonstrate that CDI-mediated interference competition actively occurs during mixed-strain infections in vivo. Together, our findings reveal that *P. aeruginosa* leverages CDI as a potent, strain-dependent kin discrimination mechanism that directly shapes population dynamics and strain dominance within host and natural environments.

**Author Summary:** Bacterial populations frequently consist of diverse strains competing for the same resources and spatial niche. We investigated how a form of interbacterial competition called contact-dependent growth inhibition (CDI) contributed to fitness and survival among diverse strains of *Pseudomonas aeruginosa*. Our findings demonstrate that bacteria deploy these contact-dependent weapons to distinguish self from non-self, governing complex competitive relationships that range from aggressive offensive killing to specialized defensive protection. Finally, using a mouse model of acute pneumonia, we confirmed that CDI actively dictates strain dominance and shapes overall bacterial burdens during infection. Together, our work shows that CDI acts as a vital mechanism for strain discrimination, directly driving population structure and strain succession in complex environments.

## Introduction

*Pseudomonas aeruginosa* is a ubiquitous environmental bacterium that occupies diverse terrestrial and aquatic niches (1). It is also an important opportunistic pathogen, frequently causing acute and chronic infections of susceptible patients (2). Within either environmental or infection contexts, communities of *P. aeruginosa* are often composed of unique conspecific strains that must continually compete for space and resources (3–6). To maintain ecological stability within these multi-strain communities, *P. aeruginosa* deploys a diverse arsenal of highly regulated interference competition mechanisms (7,8). These strategies include both contact-dependent and contact-independent weapons, such as the precision delivery of toxic effectors via its three distinct Type VI Secretion Systems (9–11) and the production of diffusible protein toxins (12–14).

Contact-dependent growth inhibition (CDI) is a specialized form of interference competition that enables many Gram-negative bacteria to directly intoxicate neighboring cells (15). CDI systems are composed of three core genes, *cdiB*, *cdiA,* and *cdiI* (**Fig. 1A**). CdiB is a TonB-dependent transporter-like outer membrane protein that displays CdiA on the outer membrane of the bacterial cell (16). CdiA is a large protein composed of multiple domains that cooperatively facilitate the delivery of its own toxic C-terminal region (CdiA-CT) into a susceptible prey cell (**Fig. 1B**) (17–21). Accordingly, CdiA-producing strains also encode a cognate immunity factor, CdiI, that binds and inactivates the specific CdiA-CT, thereby preventing self-intoxication of siblings (22–24).

**Figure 1.**
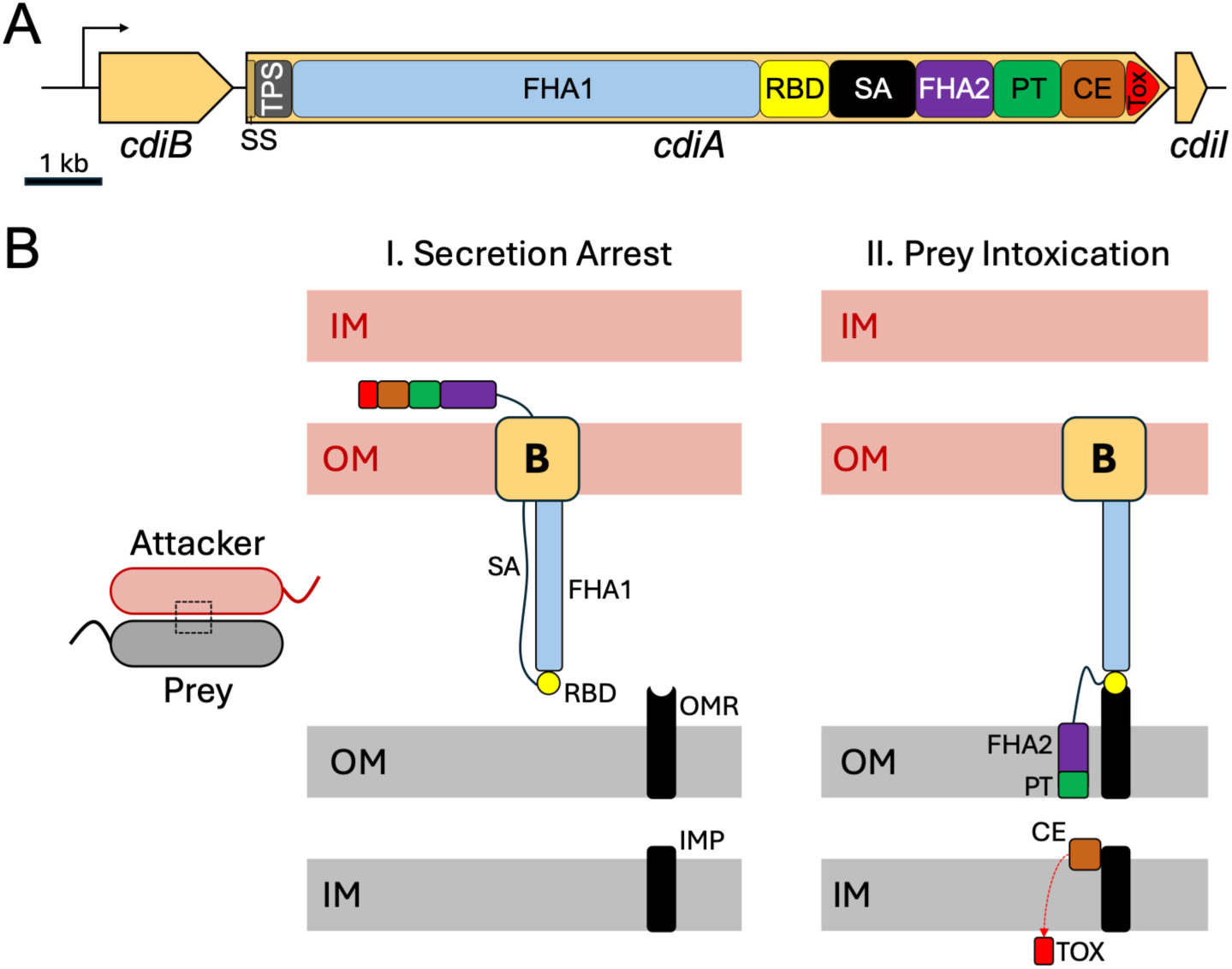
Model of contact-dependent growth inhibition (CDI). (A) Diagram of the core *cdi* locus. A single promoter drives transcription of the entire locus (arrow). CdiA protein domains are displayed as rectangles. Abbreviations are as follows: signal peptide for co-translational export via the general secretory pathway (SS), two-partner secretion domain (TPS), receptor-binding domain (RBD), secretion arrest domain (SA), pre-toxin domain (PT), C-terminal region (CT), CT cytoplasmic entry domain (CE), CT toxin domain (Tox). The FHA1 (Pfam ID PF05594) and FHA2 (Pfam ID PF13332) domains are described in the main text. (B) Current model of CdiA-mediated intoxication. Secretion of CdiA is paused (I. Secretion Arrest), with the carboxy half of the protein remaining in the attacker cell periplasm. Upon contact with the prey cell, secretion of CdiA resumes and the process of prey cell intoxication begins (Prey Intoxication). CdiA protein domains are colored according to panel B. Abbreviations include: inner membrane (IM), periplasm (PP), outer membrane (OM), outer membrane receptor (OMR), inner membrane proteins (IMP). The CdiB translocon is depicted as a black square labeled B.

CdiA export through CdiB proceeds from the amino-terminus until a specific region of CdiA is reached where secretion is halted (**Fig. 1C**) (18). In this “secretion-arrested” state, the amino half of CdiA forms a rigid filament composed of a repeating filamentous hemagglutinin peptide sequence (FHA1, Pfam ID PF05594) that displays a receptor-binding domain (RBD) distal to the cell surface. The carboxy-terminal half of CdiA remains sequestered in the periplasm of the attacking cell, linked to the RBD through a proline-rich extended region and secretion arrest (SA) domain. Binding of the RBD to a specific outer-membrane receptor (OMR) on a prey cell is a trigger that resumes CdiA export. Once released from its sequestered state, the CdiA-CT region is delivered into a prey cell through a highly coordinated, stepwise translocation process (**Fig. 1C**). First, the CdiA FHA-2 domain (Pfam ID PF13332) stably associates with the prey cell’s outer membrane, facilitating the transport of the CdiA-CT region into the periplasm through an unknown mechanism (18). During this outer membrane crossing, autoproteolytic cleavage near a conserved Val-Glu-Asn-Asn (VENN) peptide sequence releases the CdiA-CT into the periplasm of the target prey (25, 26). While some CTs function directly in the periplasm by attacking the inner membrane (27), many others target cytosolic factors and must cross the inner membrane to reach that compartment (28). This second crossing is driven by a cytoplasmic entry (CE) domain located at the N-terminus of the CdiA-CT region (**Fig. 1B**). To breach the inner membrane, specific inner membrane proteins (IMPs) in the prey cell are thought to tether the released CdiA-CT complex near the inner membrane (28, 29). IMP tethering promotes transport of the CdiA-CT toxin domain into the prey cell cytosol through an unknown mechanism (29).

CdiA is considered a polymorphic toxin because different strains can possess distinct *cdiA* and *cdiI* alleles that encode for unique CdiA-CT toxin variants and immunity proteins, respectively (30). Allelic variation within *cdiA* also occurs within the regions encoding the RBD and CdiA-CT CE domains. These different RBD and CE domain variants specifically interact with different partner proteins on the prey cell to promote intoxication. For example, CdiA-RBD variants from *E. coli* strains EC93, EC536, and STEC_O31 interact with BamA, the heterotrimeric osmoporin protein OmpC, and the outer membrane nucleoside transporter Tsx, respectively (17, 31, 32). Likewise, CdiA-CT CE variants from *E. coli* strains EC93, EC536, STEC_O31, MHI813 interact with the multidrug efflux-pump inner-membrane component AcrB, the inner membrane-anchored AAA^+^ protease FtsH, the glucose PTS permease PtsG, and ABC transporter membrane permease MetI, respectively (17, 28, 33). This target specificity in the CdiA RBD and CE domains often restricts prey cell selection at the species level (32, 34) and, along with CdiI, acts as a strict gatekeeper that broadly dictates which cells are recognized as target prey and which are embraced as cooperative kin (35). Cooperation within a microbial population is greatest when the genetic relatedness between interacting cells is high (*i.e.,* clonal or kin), such as in biofilm communities (36, 37). Bacteria can exploit CdiA’s molecular specificity to distinguish exact genetic clones from closely related prey, such as different strains within the same species (38). This physical, contact-mediated exclusion segregates mixed populations, forcing spatial clustering and establishing highly cooperative, single-strain biofilms (39).

To date, most studies on CDI have relied heavily on artificial, single-toxin chimeras overexpressed in isogenic laboratory backgrounds. While these engineered systems have been invaluable for mapping CdiA intoxication of prey, very few studies have investigated CDI-mediated competition between different strains within a single species (15, 39–41). Consequently, it remains unclear how CDI influences population dynamics independent of laboratory manipulation. Addressing this knowledge gap is critical because interference competition can influence the virulence of bacterial pathogens (42–44). In healthcare settings, genetically distinct strains of *P. aeruginosa* can maintain a permanent foothold and circulate as a highly diverse, co-colonizing pool of strains (45, 46). Consequently, patients are frequently exposed to multiple strains simultaneously, leading to complex, mixed-strain infections that not only accelerate the evolution of antibiotic resistance but also turn these shared infection sites into competitive battlegrounds (3, 47). Within these microenvironments, coexisting lineages rarely cooperate passively; instead, their genetic differences trigger fierce, direct warfare as they battle one another for survival. Determining how CdiA diversity among clinical strains shapes dominance in these polymicrobial environments is therefore vital, as it could elucidate whether CDI-mediated selection directly influences the progression and severity of patient infections.

In this study, we systematically investigated CDI during mixed-strain competitions using natural *P. aeruginosa strains*. We defined the activity of several CdiA toxin domains and mapped the competitive relationships they govern. Finally, using a mouse model of acute pneumonia, we demonstrate that CDI-mediated interference competition actively occurs during mixed-strain infections in vivo. Together, our findings reveal that *P. aeruginosa* utilizes CDI as a potent, strain-dependent kin discrimination mechanism that shapes strain dominance and population dynamics in clinically relevant host environments.

## Results

### Functional Validation and Enzymatic Diversity of *P. aeruginosa* CdiA Toxins

To investigate how CdiA toxin diversity affects competitive traits, we selected 6 phylogenetically distinct *P. aeruginosa* clinical strains from our collection that differ in their *cdiA* and *cdiI* alleles (Table 1) (48). The CdiA-CT toxin domain from PABL017 was previously determined to have tRNAse activity (49). To investigate the mechanisms of the remaining toxin alleles, we used AlphaFold3 to predict the structures of each CdiA-CT toxin domain bound to its cognate CdiI immunity factor (**Fig. S1**) (50). All CdiA-CT toxin-immunity complexes yielded high predicted template modeling (pTM) and interface predicted template modeling (ipTM) scores, confirming high-confidence models (Table 1). To gain functional insights, we searched these predicted structures against the Foldseek server (51). All toxin domains matched structural homologs with high predictive confidence, suggesting potential mechanisms of action. Most were predicted to function as nucleases, a well-established mechanism in CDI (52). The CdiA-CT toxin domain from PABL083 was predicted to be a cytidine deaminase.

**Table 1.**
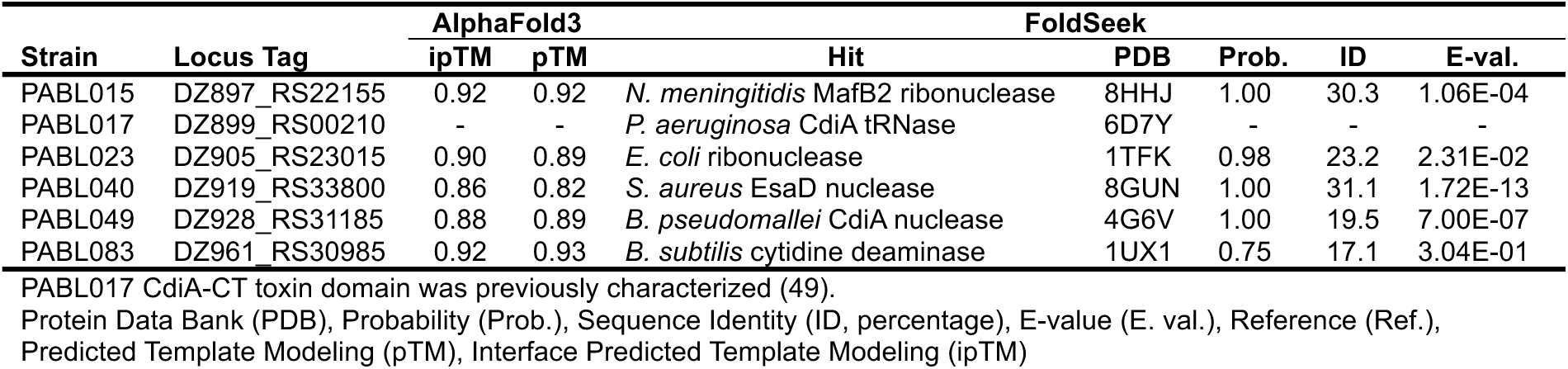
*P. aeruginosa* CdiA toxin diversity.

To confirm that the predicted CdiA-CT toxin domains impart toxic activity and that the cognate CdiI immunity proteins are protective, we cloned each pair into the pRSFDuetAra expression vector, where the *cdiA*-CT toxin domain was cloned as a functional open reading frame under transcriptional control of the arabinose-inducible araBAD promoter (*P_BAD_*) and the cognate *cdiI* was under transcriptional control of the isopropyl β-D-1-thiogalactopyranoside (IPTG)-inducible lac promoter (*P_lac_*). Each construct was transformed into *E. coli* DH5α and growth was monitored under differential inducing conditions. Arabinose induction resulted in clear growth inhibition of *E. coli,* which was partially rescued by expression of the corresponding *cdiI* immunity protein (**Fig. 2**). Thus, all cloned regions represent functional *cdiA* CT-toxin domains and their respective *cdiI* immunity protein.

**Figure 2.**
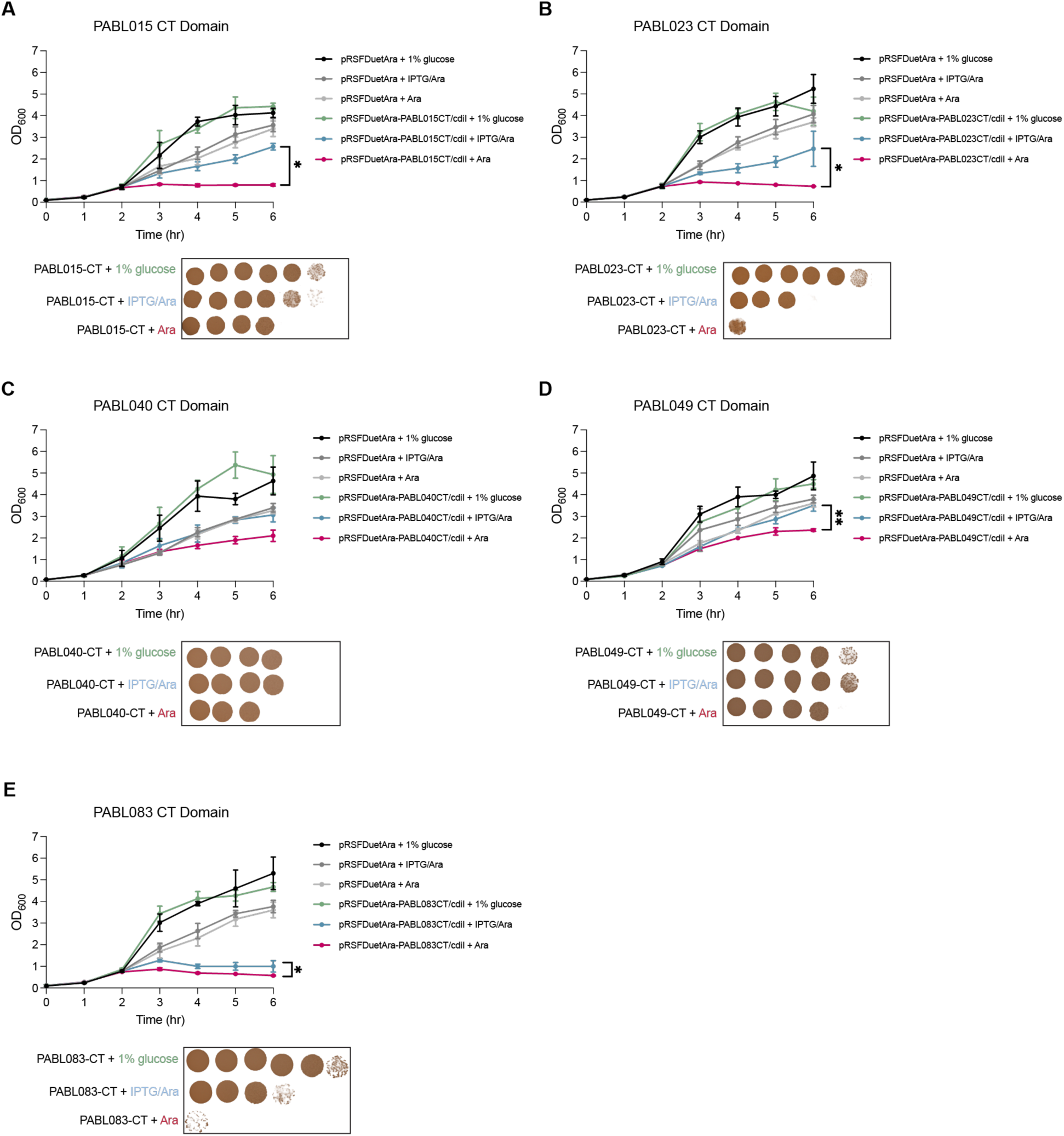
Growth inhibition of heterologously expressed CdiA CT toxin domains. *E. coli* DH5a containing either the empty vector pRSFDuetAra (black, dark grey or light grey) or vectors containing both CdiA toxin domains with their cognate immunity (green, blue, pink) from PABL015 (A), PABL023 (B), PABL040 (C), PABL049 (D), or PABL083 (E) were inoculated into LB broth containing glucose (black/green, all repressed), IPTG and Arabinose (dark grey/blue, toxin/immunity expressed), or Arabinose alone (light grey/pink, only toxin expressed). Bacterial growth was measured by monitoring OD_600_ hourly (upper panels), and viable bacterial counts were determined by plating 10-fold serial dilutions onto LB after six hours (lower panels). A two-tailed t-test was performed comparing toxin only and toxin with immunity under induction. *P < 0.05, **P < 0.01.

Because CdiA-CT toxin activity ultimately dictates CdiA-mediated competition among *P. aeruginosa* strains, we characterized the enzymatic activity of each toxin domain in vitro. Four previously uncharacterized CdiA-CT toxin domains from PABL015, PABL023, PABL040, and PABL049 were predicted to function as nucleases (**Table 1**).

After determining metal ion preferences for each variant (**Fig. S2**), purified CdiA-CT toxin domains were incubated with plasmid DNA or bulk RNA, and cleavage products were analyzed by gel electrophoresis to evaluate substrate preference (**Fig. 3**). The PABL040 toxin domain exhibited robust, concentration-dependent DNase activity with no detectable RNA cleavage (**Fig. 3C, 3G**). Conversely, PABL015 and PABL049 toxin domains displayed potent ribonuclease activity, generating specific RNA cleavage products even at the lowest concentrations tested (**Fig. 3E, 3H**); PABL049 also exhibited weak DNA cleavage at high protein concentrations (**Fig. 3D**). Finally, the PABL023 toxin domain lacked ribonuclease activity entirely (**Fig. 3F**) and yielded only minimal DNA cleavage at high protein concentrations (**Fig. 3B**).

**Figure 3.**
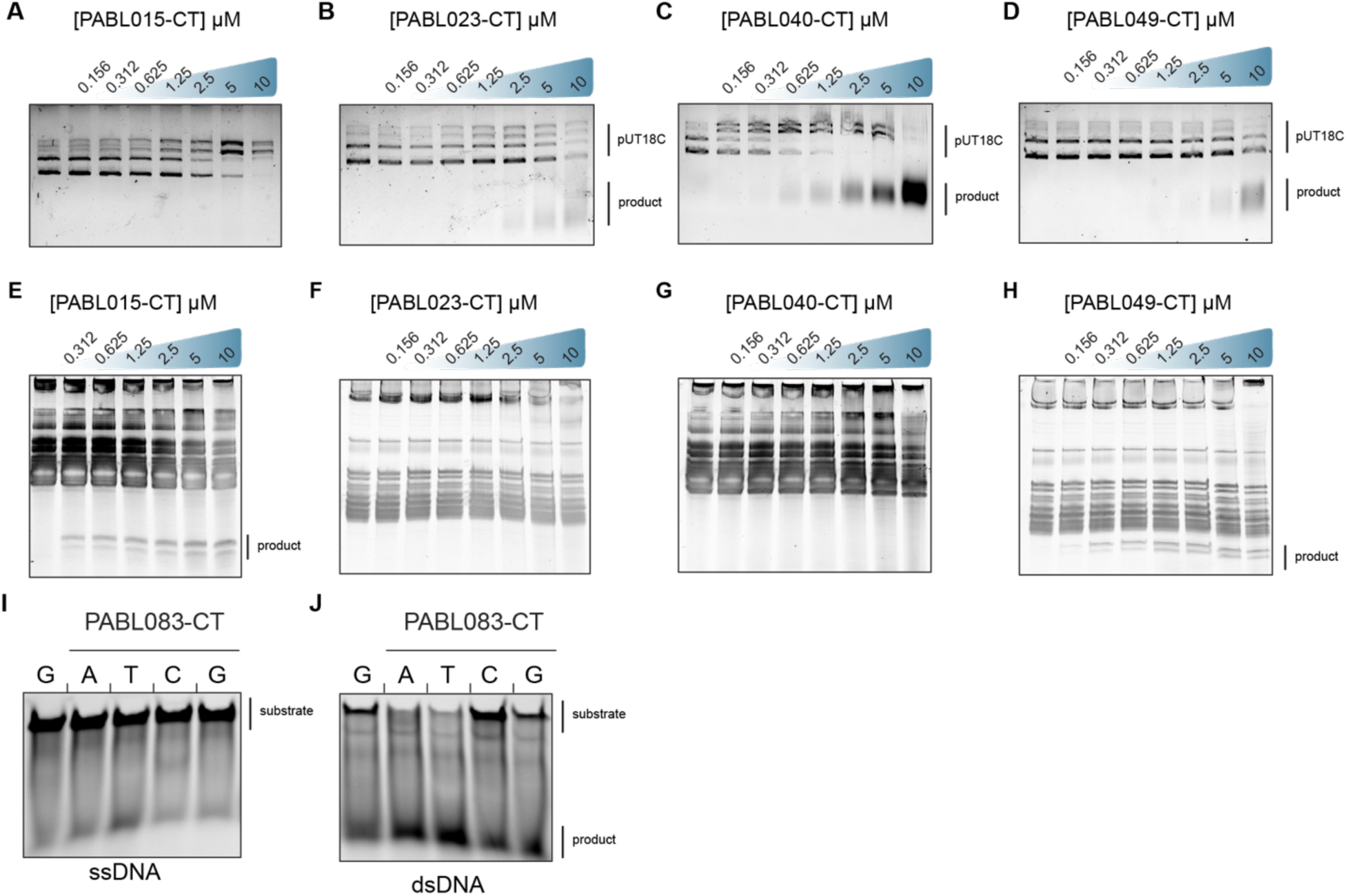
C-terminal toxin domains display diverse biochemical activities. The CdiA toxin domains (CT) from *P. aeruginosa* isolates PABL015, PABL023, PABL040, PABL049, or PABL083 were cloned, purified in isolation (without CdiI) and tested in several in vitro biochemical assays. Nuclease activity was assayed in the presence of pUT18C plasmid DNA (A-D) or *E. coli* DH5a bulk RNA (E-H) as substrates. Nuclease activity was detected by the formation of cleavage products with increasing toxin amounts, assessed by gel electrophoresis. Cytidine deamination assays for the CdiA^PABL083^ toxin domain using a single-stranded (I) or double-stranded (J) 6FAM-labeled DNA substrate with cytidines in the dinucleotide context of <u>A</u>C, <u>T</u>C, <u>C</u>C, and <u>G</u>C. Greater loss of the substrate band and stronger product bands indicate cytidine deamination of that oligonucleotide. The first lane includes a control with the GC substrate and no Toxin domain.

The CdiA toxin domain from PABL083 was predicted to function as a cytidine deaminase. Cytidine deaminases convert cytidine to uridine, triggering base excision repair (BER) (53). When BER excises closely spaced uracil lesions on opposing strands, it can cause lethal double-strand DNA breaks (54). To determine whether PABL083 possesses cytidine deamination activity, we performed in vitro deamination assays using single- and double-stranded DNA templates. The PABL083 CdiA toxin domain efficiently deaminated cytidine residues within double-stranded DNA (**Fig. 3J**), but not single-stranded DNA (**Fig. 3I**), displaying a broad sequence preference for cytidines preceded by adenosine, guanosine, or thymidine (**Fig. 3J**). Together, these results validate most of our in silico predictions and confirm that *P. aeruginosa* CdiA-CT toxin domains utilize diverse mechanisms to mediate intercellular growth inhibition.

### Orphan immunity genes provide strain-specific protection against CDI in *P. aeruginosa*

To evaluate whether candidate *cdiA* alleles confer functional CDI, we constructed CDI-susceptible, isogenic prey mutants for each strain by targeted deletion of the region encoding the CdiA-CT toxin domain and its cognate *cdiI* immunity gene (ΔCT/I). In 1:1 competition assays on solid agar, wild-type (WT) and CdiA-deficient attacker strains were co-cultured with their corresponding prey for 3 hours before quantifying prey survival. While the degree of inhibition varied, ranging from modest in PABL040 (5-fold) to dramatic in PABL083 (65-fold) and PABL017 (500-fold), nearly all WT strains significantly inhibited prey replication in a CdiA-dependent manner (**Fig. 4A**). The sole exception was PABL015, which showed no detectable CDI activity. Thus, although the CdiA-CT toxin domain of PABL015 is a functional ribonuclease (**Fig. 3E**), PABL015 does not appear to perform CDI.

**Figure 4.**
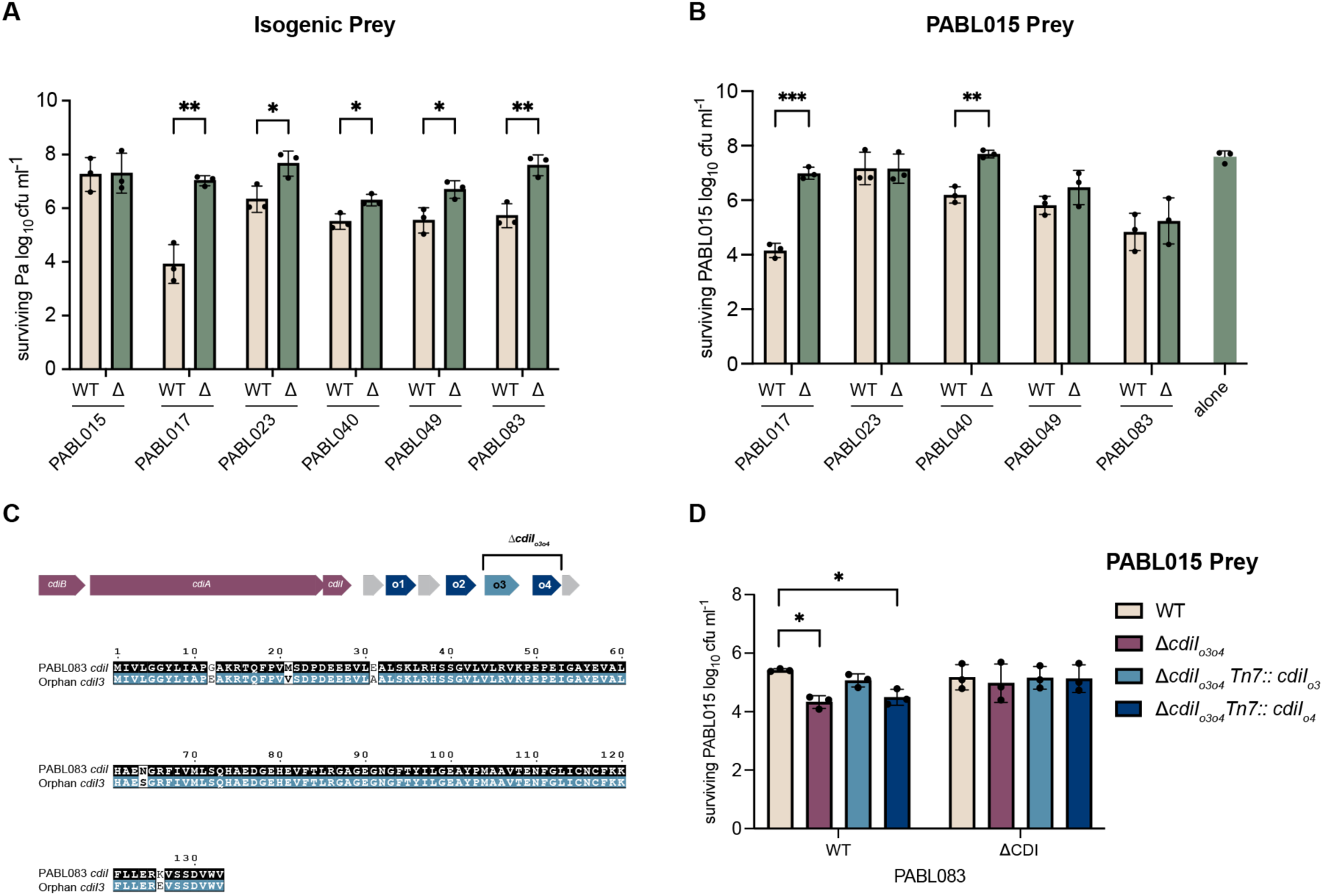
Prevalence and functional activity of contact-dependent growth inhibition systems across *P. aeruginosa* isolates. (A) Isogenic competitions. CDI-susceptible “prey” strains missing the toxin C-terminal domain and immunity gene (ΔCT/I) were co-cultured at a 1:1 ratio with wild-type or ΔCT/I attacker strains on solid media for 3 hr at 37 °C. Surviving prey cells were quantified by serial dilution plating on selective media. Strains are indicated on the x-axis. Statistical significance was evaluated using multiple unpaired t-tests with Holm–Šídák correction. (B) Mixed-strain competitions between isolate PABL015 (prey) and indicated *P. aeruginosa* attacker strains (WT tan; or ΔCT/I, green) were performed as described in panel A. (C) Schematic of the PABL015 *cdi* locus highlighting four orphan *cdiI* immunity genes (*cdiI*_o1_–*cdiI*_o4_), the double mutant construct (Δ*cdiI*_o3o4_), and an amino acid sequence alignment between *cdiI*_o3_ and *cdiI* from isolate PABL083. (D) Mixed-strain competitions evaluating protection by orphan immunity genes. Wild-type PABL015 (tan), Δ*cdiI*_o3o4_ (pink), and Δ*cdiI*_o3o4_ complemented at the Tn7 locus with orphan *cdiI*_o3_ (light blue) or *cdiI*_o4_ (dark blue) were competed against PABL083 attacker strains (WT, tan; or ΔCT/I, green). Statistical significance between prey strains was evaluated using a two-way ANOVA with Tukey’s multiple comparisons test. Data across all panels represent the mean ± SD from three independent experiments performed in triplicate. ∗P<0.05, P<0.01,∗∗∗P<0.001.

The observation that distinct *P. aeruginosa* strains execute CDI to varying degrees highlights CdiA’s potential to promote strain dominance. We therefore investigated mixed-strain CDI as a potential mechanism for kin discrimination in *P. aeruginosa*. Because PABL015 lacks CDI activity, we selected it as a universal prey strain to evaluate attacker activity without the confounding variable of reciprocal killing. Using this setup, pairwise competition assays were conducted with PABL017, PABL023, PABL040, PABL049, and PABL083 as potential attacker strains. Among the six candidates, strains PABL017 and PABL040 exhibited significant CdiA-dependent killing against PABL015 (**Fig. 4B**). In contrast, PABL023, PABL049, and PABL083 did not exhibit *cdiA*-dependent killing, although these isolates were able to inhibit PABL015 independent of *cdiA*. In many bacteria, *cdi* loci harbor genomic sequences displaced from full-length *cdiA* and *cdiI* genes (55). Orphan *cdiI* genes have previously been shown to confer protection against competitors harboring matching CdiA toxins (55).

Genomic analysis downstream of *cdiBAI* in strain PABL015 revealed four orphan toxin-immunity modules, two of which correspond to the active CdiA-CT toxin domains and cognate CdiI immunity factors encoded by PABL083 and PABL023 (**Fig. 4C, Fig. S3A**). To test whether these orphan *cdiI* genes protected PABL015, we deleted both genes (*cdiI*_o3_ and *cdiI*_o4_) from PABL015 and assayed the mutant in competition against PABL023 and PABL083. Deletion of the orphan immunity genes enabled PABL083 to execute CDI-mediated killing of PABL015 (**Fig. 4D**). Protection against PABL083 was successfully restored by complementation with the cognate *cdiI*_o3_ gene, but not the non-cognate *cdiI*_o4_ gene (**Fig. 4D**), confirming that orphan immunity genes naturally protect against CdiA-mediated killing by competing isolates with partner toxin alleles. In contrast, PABL023 was still unable to demonstrate CdiA-mediated killing against the PABL015 Δ*cdiI*_o3o4_ strain (**Fig. S3D**). These results indicate that the lack of mixed-strain CDI activity in PABL023 stems from mechanisms outside of orphan immunity genes.

Together, these findings show that orphan immunity genes can mediate cross-strain defense against specific CDI systems, while additional, uncharacterized barriers can shape the overall outcomes of mixed-strain competition.

### CdiA toxin production shapes, but does not dominate, competitive outcomes among *P. aeruginosa* isolates

We expanded our pairwise competitions to the remaining CDI-competent strains, and characterized the impact of CdiA on mixed-strain outcomes across all four possible genotype combinations per pairing (*i.e*., WT vs. WT, WT vs. Δ, Δ vs. WT, and Δ vs. Δ). Surviving cells at 3 hours were quantified using selective plating to distinguish each competitor, and a competitive index (CI) was calculated to determine the strain/genotype with the competitive advantage (i.e., winner) or competitive disadvantage (i.e., loser) (**Table 2**). Rather than acting purely as offensive weapons, CdiA-mediated systems served diverse functional roles across strains, ranging from offensive aggression to defensive protection.

**Table 2.** Outcome of pairwise competition assays across genotype combinations.

| Pair | WT : WT | $\Delta$ : WT | WT : $\Delta$ | $\Delta$ : $\Delta$ | CDI Influence |
| --- | --- | --- | --- | --- | --- |
| 17 vs. 23 | -0.07 ($\pm 0.54$)<br>PABL023 | -5.31 ($\pm 0.43$)<br>PABL023 | 2.31 ($\pm 0.81$)<br>PABL017 | -3.73 ($\pm 0.70$)<br>PABL023 | Determines winner |
| 17 vs. 40 | 0.34 ($\pm 0.15$)<br>PABL017 | 0.59 ($\pm 0.35$)<br>PABL017 | 2.8 ($\pm 0.07$)<br>PABL017 | 0.93 ($\pm 0.35$)<br>PABL017 | Amplifies severity of loss |
| 17 vs. 49 | -1.70 ($\pm 0.62$)<br>PABL049 | -1.85 ($\pm 0.87$)<br>PABL049 | -0.65 ($\pm 0.97$)<br>PABL049 | -0.71 ($\pm 0.74$)<br>PABL049 | Amplifies severity of loss |
| 83 vs. 17 | 5.08 ($\pm 0.07$)<br>PABL083 | 1.18 ($\pm 0.25$)<br>PABL083 | 8.22 ($\pm 0.28$)<br>PABL083 | 5.61 ($\pm 0.35$)<br>PABL083 | Amplifies severity of loss |
| 83 vs. 23 | 1.86 ($\pm 0.19$)<br>PABL083 | 1.45 ($\pm 0.34$)<br>PABL083 | 2.55 ($\pm 0.22$)<br>PABL083 | 1.71 ($\pm 0.03$)<br>PABL083 | Amplifies severity of loss |
| 83 vs. 40 | 7.11 ($\pm 0.08$)<br>PABL083 | 0.17 ($\pm 0.07$)<br>PABL083 | 7.31 ($\pm 0.11$)<br>PABL083 | 0.05 ($\pm 0.09$)<br>PABL083 | Amplifies severity of loss |
| 83 vs. 49 | 0.11 ($\pm 0.54$)<br>PABL083 | -1.95 ($\pm 0.77$)<br>PABL049 | 1.41 ($\pm 0.69$)<br>PABL083 | -0.18 ($\pm 0.14$)<br>PABL049 | Determines winner |
| 83 vs. 17<br>(1 hr) | 3.21 ($\pm 0.56$)<br>PABL083 | 0.06 ($\pm 0.05$)<br>PABL083 | 5.40 ($\pm 1.20$)<br>PABL083 | 0.01 ($\pm 0.06$)<br>PABL083 | |
| 83 vs. 40<br>(1 hr) | 3.56 ($\pm 0.93$)<br>PABL083 | -0.14 ($\pm 0.10$)<br>PABL040 | 4.28 ($\pm 0.33$)<br>PABL083 | -0.13 ($\pm 0.19$)<br>PABL040 | |

Cell entries show average Log10 competitive index (CI) from three independent biological replicates with standard deviation (n = 3 replicates per condition). Three-hour competitions unless otherwise noted. Negative Log10 CI indicates the 2<sup>nd</sup> strain in the pair wins. Winning strain is noted underneath values.

In five of the seven pairings (PABL017 vs. PABL040, PABL017 vs. PABL049, PABL083 vs. PABL017, PABL083 vs. PABL023, and PABL040 vs. PABL083), the same strain prevailed regardless of whether *cdiA* was intact (**Table 2**), indicating that the dominant strain did not require CdiA to outcompete its rival. CdiA nonetheless shaped the magnitude of these competitions through distinct pairing-specific mechanisms.

In two of these five pairings, CdiA had only a minor effect on the margin of victory. Mutating *cdiA* in PABL083 slightly reduced its advantage over PABL023, whereas PABL023’s *cdiA* genotype had no significant impact on the outcome (**Fig. 5D**). Similarly, CdiA contributed marginally to PABL049’s victory margin, becoming significant only when competing against the PABL017 *cdiA* mutant (**Fig. 5B**).

**Figure 5.**
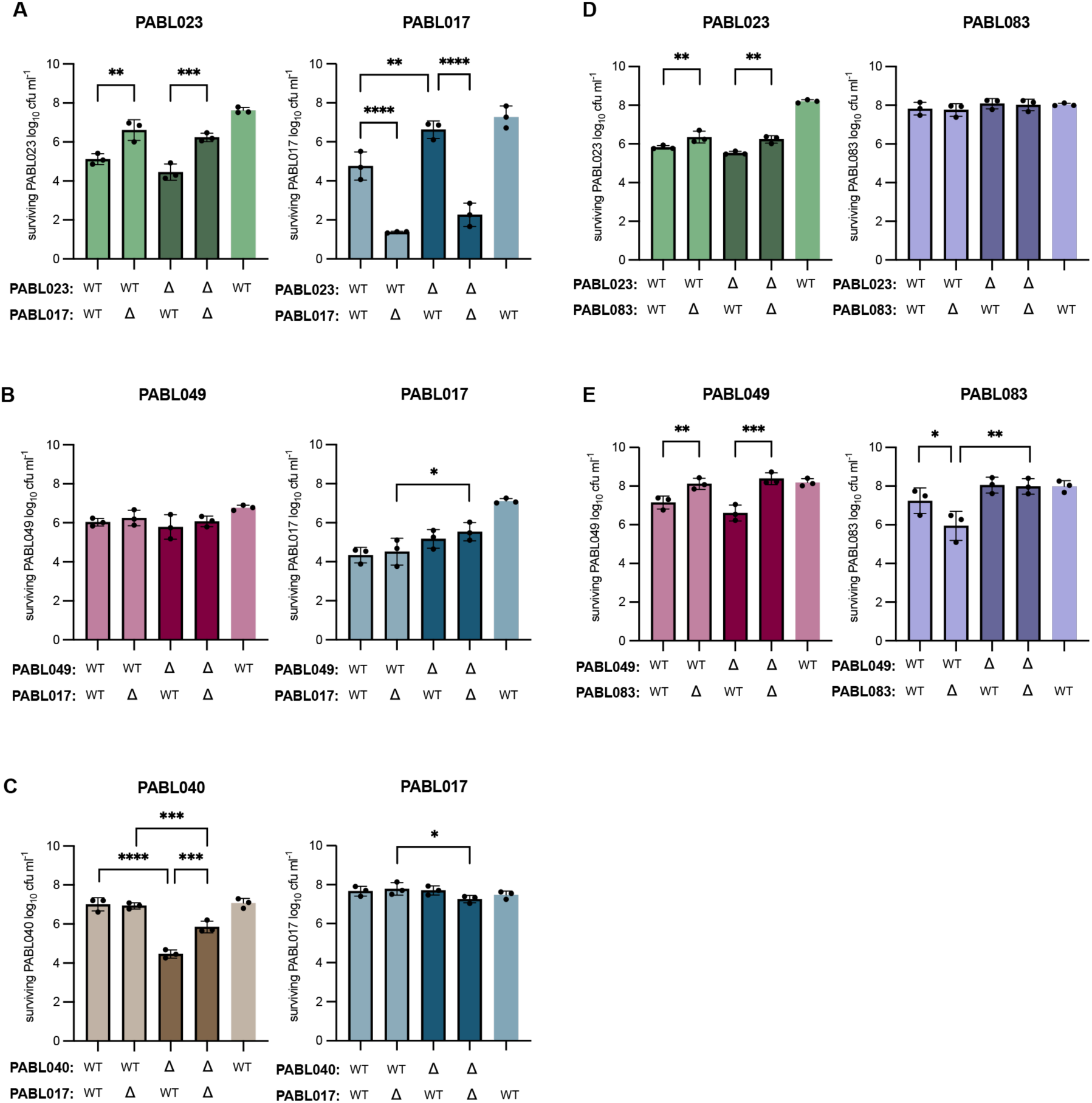
Mixed-strain competitions. (A) Mixed-strain competitions were performed by co-culturing isolate A WT or Δ*cdiA*-Tox Δ*cdiI* (Δ) strains at a 1:1 ratio with the respective strains from isolate B, and incubating on solid media for 3 hr. at 37 °C. The initial bacterial numbers at the start of competition (Input) and the surviving bacteria after competition (3 hr,) were quantified by plating serial dilutions on selective media. Data represent the mean ± SD from three independent experiments performed in triplicate. Bars are colored according to isolate: PABL017 (blue), PABL023 (green), PABL040 (tan), PABL049 (red), and PABL083 (purple). Strain pairs evaluated in each mixed competition are indicated on the x-axis. Data represent the mean ± SD from three independent experiments performed in triplicate. Statistical significance between different strains was evaluated using a Two-Way ANOVA with Tukey’s multiple comparison test (**P<0.01, ***P<0.001).

In contrast, CdiA played a far greater role in defining the competitive outcome in the remaining three pairings. For PABL017 vs. PABL040, the losing strain’s genotype governed the interaction. PABL017 dramatically defeated PABL040 when PABL040 lacked *cdiA* (Log10 CI = 2.4), but only minimally across all other genotype combinations (**Fig. 5C**). This demonstrates that PABL040 deploys CdiA defensively to mitigate population collapse caused by PABL017. A more dramatic pattern emerged in the PABL040 vs. PABL083 pairing, where PABL040 was nearly eliminated whenever PABL083 possessed functional CdiA, regardless of PABL040’s own genotype (Log10 CI > 7.1) (**Fig. 6B**). This killing was completely dependent on PABL083 encoding *cdiA*, as PABL083 *cdiA* mutant strains showed no competitive advantage (Log10 CI < 0.17). Suppression began within 1 hour, confirming rapid kinetic onset of CdiA-dependent killing by PABL083 (**Fig. 6D**).

**Figure 6.**
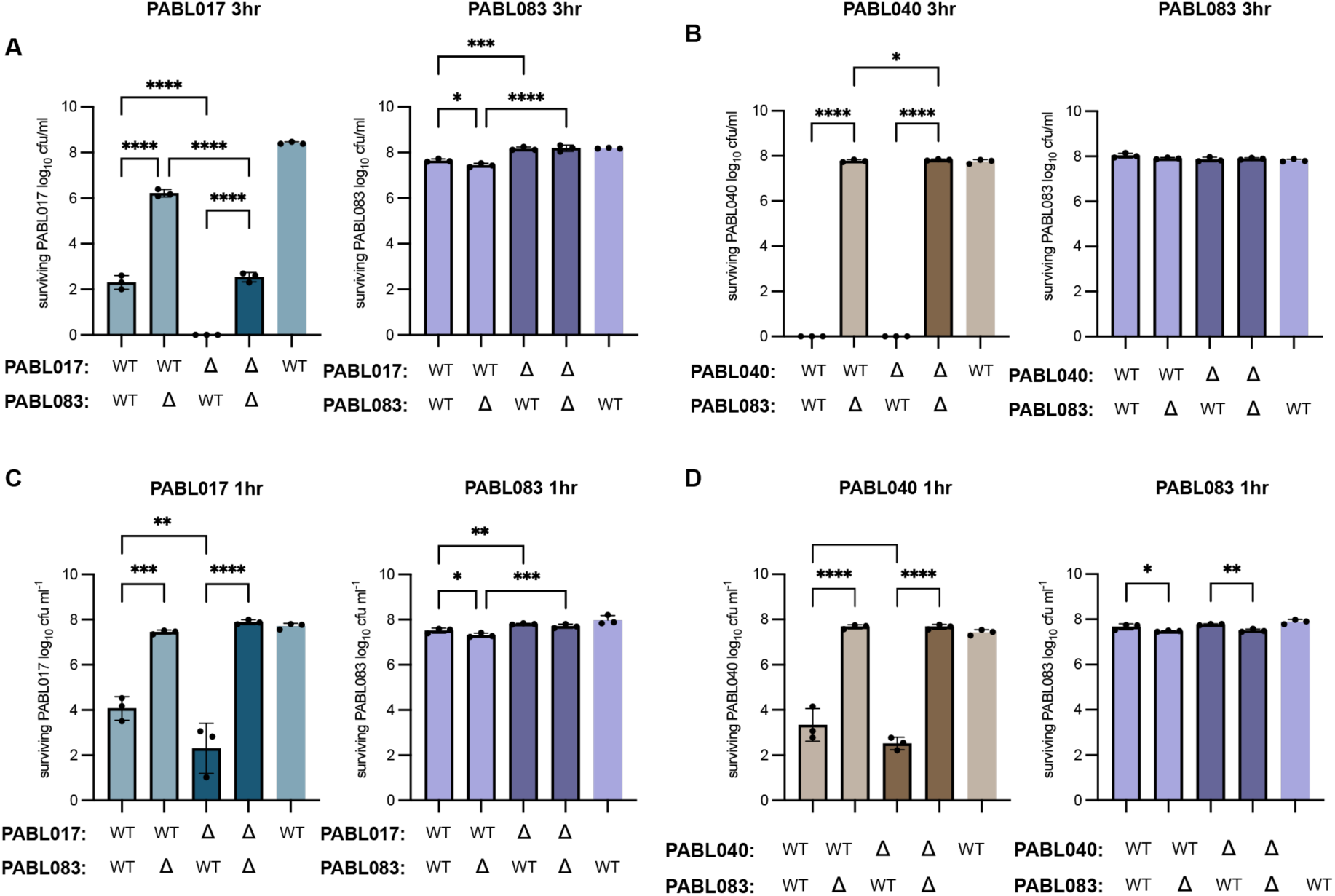
PABL083 mixed-strain competitions. (A) Mixed-strain competitions were performed as described in Figure 5, comparing PABL083 strains with the respective PABL017 (A, C) or PABL040 (B, D) strains. The surviving bacteria after 3 hr. competition (A, B) or 1 hr. competition (C, D) were quantified by plating serial dilutions on selective media. Data represent the mean ± SD from three independent experiments performed in triplicate. Bars are colored according to isolate: PABL017 (blue), PABL040 (tan), or PABL083 (purple). Strain pairs evaluated in each mixed competition are indicated on the x-axis. Data represent the mean ± SD from three independent experiments performed in triplicate. Statistical significance between different strains was evaluated using a Two-Way ANOVA with Tukey’s multiple comparison test (*P<0.05, **P<0.01, ***P<0.001, ****P<0.0001).

Finally, PABL083 and PABL017 engage in a distinct competitive interplay driven by two complementary mechanisms. PABL083 strongly inhibited PABL017 (Log10 CI = 5.08). This advantage was largely lost in a PABL083 *cdiA* mutant (Log10 CI = 1.18) (**Fig. 6A**). Conversely, PABL083’s dominance increased significantly in competition with a PABL017 *cdiA* mutant (Log10 CI = 8.22), suggesting PABL017 uses its own CdiA defensively in this pairing. However, when both strains were *cdiA* mutants, PABL083 still outcompeted PABL017 comparable to WT competition (Log10 CI = 5.61), suggesting that PABL083 has additional CdiA-independent competitive mechanisms. Shorter experiments show that CdiA-mediated inhibition acts rapidly, suppressing PABL017 within the first hour of co-culture (**Fig. 6C**). In contrast, the CdiA-independent competitive mechanism was not yet evident at this early time point. This indicates that the CdiA-independent competitive mechanism does not impact PABL017 until between 1 and 3 hours of co-culture. Thus, PABL083 relies on a biphasic strategy: fast-acting CdiA strikes immediately to establish early dominance, followed by a secondary, CdiA-independent mechanism that engages later to complete competitive clearance.

In contrast to the unconditional dominance competition patterns described above, two pairings (PABL017 vs. PABL023 and PABL049 vs. PABL083) revealed that CdiA helped dictate the winner (**Table 2**). PABL049 and PABL083 showed mutual offensive escalation, with each strain deploying CdiA toxins against the other. Both isolates were evenly matched when both encoded functional CdiA toxins (PABL083 Log10 CI = 0.11) or when both lacked them (PABL049 Log10 CI = 0.18); however, deleting either strain’s CdiA-CT toxin domain caused a severe competitive loss giving the competitive advantage to the WT strain (Log10 CI > 1.4) (**Fig. 5E**). Conversely, PABL017 and PABL023 were closely matched when both contained CdiA toxins (PABL023 Log10 CI = 0.07), but each lost when its CdiA-CT toxin domain was mutated (**Fig. 5A**). PABL023 also displayed a CdiA-independent competitive mechanism over PABL017 when CdiA-CT toxin domain was mutated from both strains (PABL023 Log10 CI = 3.73), like PABL083. Thus, whether a CdiA-CT toxin dictates competitive success in this pairing ultimately depends on the genotypic landscape of the competitor.

### CDI Shapes Inter-Strain Competition In Vivo

The finding that CDI acts as a mechanism of kin discrimination that shapes inter-strain interactions in vitro highlights the potential importance of these interactions during mixed-strain infection. To evaluate these interactions in vivo, we selected PABL017 and PABL083 because this isolate pairing had a strong in vitro competition phenotype. Mice were infected in a pneumonia model with an equal ratio of the same PABL017 and PABL083 strain combinations used above. At 5 hours post-infection, PABL083 outcompeted PABL017 in the lung by 100-fold (**Fig. 7A**). Consistent with our in vitro findings, this competition was abolished in the lung when in competition with the PABL083 *cdiA* mutant strain. However, unlike the in vitro conditions, PABL017 did not show significant CdiA-dependent killing against PABL083 in the lung (**Fig. 7A**).

**Figure 7.**
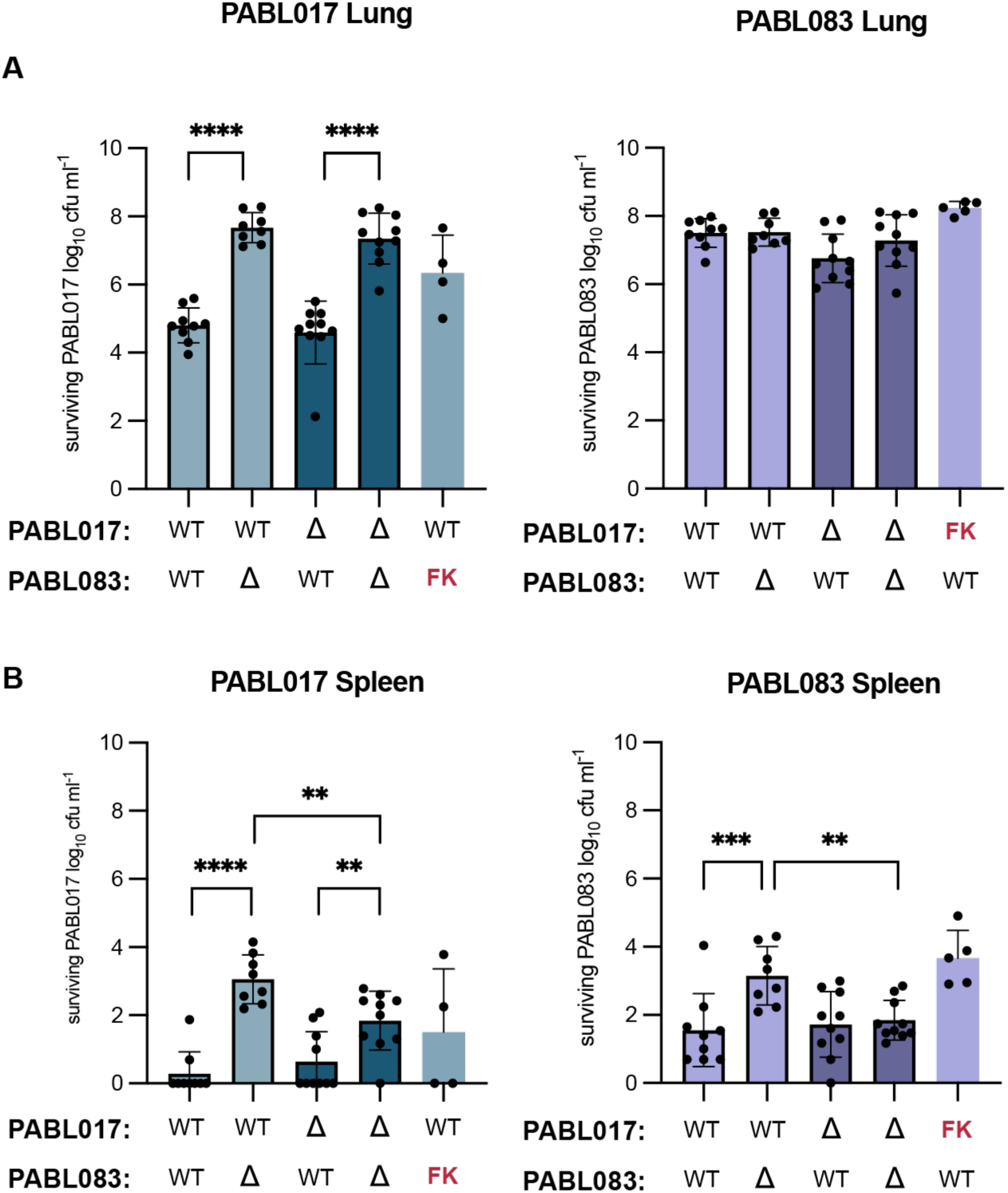
CDI of *P. aeruginosa* strains in a murine pulmonary infection model. Mixed-strain infections were initiated by intranasal co-inoculation of PABL017 wild-type (WT) or Δ*cdiA*-CT Δ*cdiI* (Δ) strains at a 1:1 ratio with corresponding PABL083 strains. At 5 hours post-infection, lungs (A) and spleens (B) were harvested, homogenized, and serial dilutions were plated on selective media to quantify surviving bacterial burdens (CFU/organ). Bars represent isolate identity: PABL017 (blue) and PABL083 (purple). Specific co-inoculated strain combinations are indicated on the x-axis. Data are presented as mean ± SD. Statistical significance was determined by two-way ANOVA with Tukey’s post-hoc multiple comparison test ( *P<0.05, **P<0.01, *** P<0.001). Abbreviations: FK: Formalin-killed

We also investigated the impact of competitor viability on lung colonization by co-inoculating live WT strains with formalin-killed (FK) competitors. Recoverable PABL083 counts in the lung did not differ whether PABL017 was viable or FK; however, PABL017 counts were reduced when mixed with viable PABL083 compared with FK PABL083, supporting the idea that PABL083 actively suppresses PABL017 numbers in the lung.

We next examined whether CDI-mediated competition influences bacterial dissemination from the lung by measuring bacterial burdens in the spleen (**Fig. 7B**). Bacterial burdens in the spleen generally mirrored lung burdens; when total burdens in the lung were higher, burdens in the spleen were correspondingly higher. This suggests that CDI affects systemic dissemination by impacting local bacterial density in the lung. The exception to this pattern occurred when both strains did not produce a functional CdiA toxin (Δ vs Δ). In this pairing, total lung burdens were high, but spleen burdens were significantly lower than in co-infections where PABL017 retained a functional CdiA toxin (WT vs Δ). This divergence suggests that the PABL017 CdiA toxin may modestly enhance dissemination independently of interbacterial competition. Together, these results demonstrate that CDI-mediated inter-strain interactions shape competitive dominance within the lung and partially impact systemic dissemination to the spleen.

## Discussion

By comparing natural *P. aeruginosa* isolates, this study addresses a gap noted throughout the CDI literature: whether *cdiA* diversity translates into predictable competitive outcomes among naturally co-circulating strains, and whether it matters in relevant contexts such as infection (35, 38, 48). Our results show that they do, but with a degree of context-dependence that is difficult to appreciate from laboratory-engineered systems alone.

The functional characterization of the CdiA toxin domains adds to a growing appreciation that CDI effectors are mechanistically diverse. Nuclease activity remains the most common mechanism in CDI and related polymorphic toxin systems (30, 52). However, the discovery that PABL083 functions as a cytidine deaminase extends this repertoire. Deaminase toxins have only recently been described as interbacterial weapons for a Type VI secretion system in *Burkholderia cenocepacia* (56). Our findings suggest that distinct contact-dependent delivery platforms may have convergently recruited the same enzymatic strategy. Together with a recently described family of membrane-disrupting CDI ionophores in *E. coli* (57), these findings suggest that continued surveys of natural CdiA diversity will likely uncover further mechanisms of interbacterial antagonism beyond the handful characterized to date.

The observation that the CdiA toxin domain of PABL023 displays inhibitory activity when expressed in *E. coli* (**Fig. 2B**) and in natural competition experiments (**Fig. 4A**), but not during in vitro nuclease experiments (**Fig. 3B, 3F**), suggests that either the experimental conditions were inappropriate, our functional predictions were incorrect, or that the PABL023 CdiA toxin domain requires a host-encoded permissive factor for activity. Several CDI tRNase toxins require host-encoded permissive factors, such as CysK in uropathogenic *E. coli* 536 or elongation factor Ts in enterohemorrhagic *E. coli* EC869, to achieve full activity in the target cytosol (58, 59). The failure of PABL023’s toxin domain to cleave nucleic acids efficiently in vitro, despite confident structural prediction of nuclease activity, is therefore not unprecedented and will require further work to experimentally validate our functional predictions.

The role of orphan immunity modules in protecting PABL015 against CDI by PABL083 highlights a unique attribute of CDI evolution (**Fig. 4C**). First described in *Burkholderia* and enteric systems, truncated orphan *cdiA-cdiI* fragments retained in the genome provide immunity against toxins encoded by full-length systems elsewhere in the population (55). Orphan immunity genes linked to other polymorphic toxin delivery platforms, most extensively the Type VI secretion system (T6SS), are well established as a defense strategy: orphan Rhs immunity alleles protect *Salmonella* against toxins deployed by evolved, T6SS-competent siblings (60), horizontally acquired orphan immunity gene clusters protect Bacteroidales species against T6SS attack both in vitro and within the gut of gnotobiotic mice (61), and structural work has shown that orphan immunity proteins can neutralize T6SS nuclease effectors through the same conformational mechanism used by their cognate counterparts (62). In each of these cases, however, protection was demonstrated for T6SS, and largely in the context of engineered or evolved laboratory strains rather than direct competition between unmodified, naturally occurring isolates. That two of PABL015’s four orphan modules corresponded to active toxins from other isolates in this collection, and that deleting the matching orphan immunity gene restored susceptibility to CDI during head-to-head competition (**Fig. 4D**), extends this defense strategy to CDI. To our knowledge, this provides the first demonstration that orphan immunity genes can protect against a competitor’s toxin during competition with natural strains. This supports the narrative that orphan modules are not simply evolutionary relics but can function as an operational layer of strain-specific defense under conditions that closely approximate natural polymicrobial competition, effectively giving a strain partial immunological memory of toxins it does not itself produce. This complements the broader model of CDI as a kin- and kind-discrimination system (36, 38, 39): orphan immunity may allow a strain to tolerate the presence of certain non-self-genotypes it would otherwise be vulnerable to, subtly reshaping which strains can coexist even in the absence of reciprocal toxin production.

Perhaps the most striking outcome of the pairwise competition assays is the diversity of roles that CdiA played across isolate pairs, ranging from offensive to defensive. This context-dependence echoes findings from other contact-dependent weapon systems. In *P. aeruginosa* and other species, the Type VI secretion system can be deployed either offensively, firing constitutively at neighboring cells, or defensively, assembling only in response to a detected incoming attack, a strategy formalized as bacterial "tit-for-tat" retaliation (63, 64). The behavior observed here for PABL040 and, in a more conditional form, PABL017, in which CdiA activity limited losses against a specific competitor’s toxin but conferred little independent offensive advantage, is conceptually analogous to a defensive or retaliatory weapon strategy, even though CDI, unlike the T6SS, is not known to sense incoming attack directly. Whether the defensive benefit of CdiA in these pairings reflects reciprocal toxin exchange, competition for the same outer-membrane receptor, or another mechanism remains an open question.

Conversely, the mutual escalation observed between PABL049 and PABL083, in which each isolate’s competitiveness collapsed when its own toxin was deleted regardless of the opponent’s genotype, is more consistent with an arms-race dynamic in which both lineages have come to depend on CdiA to maintain competitive parity, a pattern that aligns with theoretical and empirical work describing polymorphic toxin systems as products of continual coevolutionary escalation between competing lineages (30, 35).

This functional diversity in CdiA deployment also aligns with broader theoretical and empirical principles governing microbial warfare. A recent agent-based modeling and experimental study comparing contact-dependent weapons with long-range, diffusible weapons (such as pyocins) showed that contact weapons remain effective even when the attacking strain is in numerical minority, whereas diffusible weapons require local abundance to reach inhibitory concentrations (35). While not directly tested here, CDI may be critical for genotypes invading or persisting within established, numerically dominant populations, a scenario directly relevant to chronic airway co-colonization.

Alternatively, CDI has been documented to mediate microcolony exclusion (39, 65), and similar border-defense or retaliatory mechanisms have been documented for T6SS deployment (63, 64) and *Bacillus subtilis* biofilm exclusion systems (66, 67). In this light, defensive CdiA deployment reflects a recurring evolutionary strategy where a disadvantaged lineage uses toxin production to raise the cost of encroachment and maintain territory against aggressive competitors.

In the murine pneumonia model, CDI-mediated competition directly shaped infection outcomes, mirroring the in vitro dominance of PABL083 and its loss upon *cdiA* inactivation, confirming that direct interbacterial antagonism operates in vivo (68, 69). Conversely, PABL017’s defensive CdiA benefit was lost in the lung, indicating that spatial structure, cell density, or physiological state within host tissues can alter CDI efficacy. Notably, CDI-driven strain dominance correlated with bacterial burdens in both the lung and spleen, suggesting that local interbacterial competition influences systemic dissemination. These findings carry important clinical implications. Genomically diverse *P. aeruginosa* isolates frequently co-colonize patients (45, 46) and complicate treatment trajectories (3, 47); our data show that CDI-mediated interference competition is a primary force determining strain dominance and replacement.

Future work in this field should prioritize identifying the CdiA-independent inhibitory mechanisms evident in several pairings, testing whether the secondary mechanism in the biphasic PABL017-PABL083 interaction is a diffusible weapon such as a pyocin with distinct density-dependent behavior (35). Expanding in vivo testing to additional isolate pairs will clarify how host environments reshape diverse CDI dynamics. Determining whether *cdiBAI* expression is constitutive or regulated by quorum or competition sensing (70), and whether single cells display heterogeneous offensive or defensive strategies (71), will elucidate the regulatory basis for these behaviors. Finally, testing orphan immunity gene content across larger clinical cohorts will help extend the kin- and kind-discrimination framework (36, 38, 39, 55) toward a predictive understanding of polymicrobial *P. aeruginosa* infection.

## Materials and Methods

### Bacterial Strains and Growth Conditions

All bacterial strains, plasmids, and oligos used in this study are listed in Tables S1 through S3. Bacteria were routinely cultured in Lysogeny Broth (LB) at 37°C with aeration at 225 rpm or on agar plates. Antibiotics were supplemented as required at the following concentrations for *E. coli* or *P. aeruginosa*, respectively: gentamicin (Gm), 30 or 100 µg/mL; carbenicillin (Carb), 50 or 500 µg/mL. Irgasan (Irg), 5 µg/mL, was used to select for *P. aeruginosa*.

### Plasmid Construction

To generate the base pRSFDuetAra plasmid, the arabinose-inducible promoter from pSB109 was amplified by PCR and Rolling Circle Amplification assembled into digested pRSFDuet using Phusion High-Fidelity DNA Polymerase (Thermo Fisher Scientific).

To generate the pRSFDuetAra CT domain/immunity dual expression plasmids, sequences were amplified from *P. aeruginosa* (PABL015, PABL023, PABL040, PABL049, PABL083, PABL098) genomic DNA by PCR. Immunity PCR products were Gibson assembled into pRSFDuetAra digested with NcoI and EcoRI for insertion of the immunity (NEB). CT domain PCR products were Gibson assembled (NEB) according to the manufacturers protocol into pRSFDuetAra-immunity digested with NdeI and XhoI for insertion of the CT domain (NEB).

To generate pE-SUMO-CT domain/immunity plasmids for protein expression and purification, sequences were amplified from *P. aeruginosa* (PABL015, PABL023, PABL040, PABL049, PABL083, PABL098) genomic DNA by PCR. PCR products were Gibson assembled into BsaI-digested pE-SUMO. All oligonucleotides were purchased from IDT or Thermo Fisher Scientific.

### Generation of *P. aeruginosa* Mutants

Markerless deletions were generated using the pEXG2 suicide vector as previously described (72, 73). Briefly, 750-500 bases up and downstream of the locus to be removed were PCR amplified with upstream and downstream forward and reverse primers. Upstream reverse and downstream forward primers were designed with complementary overlap regions to allow for overlap extension PCR or Gibson assembly into digested (EcoRI/BamHI) pEXG2. Mutagenesis plasmids were maintained in *E. coli* NEBα or DH5α, then transformed into *E. coli* SM10 for conjugation. Briefly, *P. aeruginosa* was streaked onto LB and after overnight growth at 37°C, incubated at 42°C for 2 hours to increase conjugation efficiency. A sweep of fresh SM10 and *P. aeruginosa* colonies were mixed, spotted on an LB plate, then incubated at 37°C for 4 hours. The spot was resuspended and plated onto selective LB (gentamicin 100 μg/mL, irgasan 5 μg/mL) media for single colonies. Gentamicin-resistant colonies were outgrown in LB without NaCl for 3 hours and streaked onto LB supplemented with 15% sucrose plates for counterselection. Colonies were screened for antibiotic sensitivity, by PCR, and deletions confirmed by amplicon sequencing.

### Bacterial Growth Curves

After overnight growth in LB broth containing antibiotic and 1% glucose (to suppress CT domain expression), *E. coli* were normalized to an OD600 of 0.1 in LB broth containing antibiotics, 1% glucose, and 1mM IPTG. Cultures were incubated at 37°C with shaking for 1 hour, pelleted, washed with PBS, and resuspended in LB broth supplemented with either 0.2% arabinose (for induction of CT domains alone) or 0.2% arabinose and 1 mM IPTG (for induction of CT domains and cognate immunity).

Bacterial growth and viability were determined by measuring the OD600 every hour for 6 hours and by measuring CFU at 6 hours by plating 10-fold serial dilutions on LB agar containing 1% glucose, 1mM IPTG, and antibiotics.

### Bacterial Competition Assays

Overnight cultures of *P. aeruginosa* were diluted 1:50 into fresh LB supplemented with antibiotic (when appropriate) and grown to mid-exponential phase. Cells were pelleted, normalized to an OD600 of 1.0, and mixed in equal ratios. Competition mixtures were spotted as 5-10 μL drops onto LB agar and incubated for 1-3 hours at 37°C. After incubation, competition spots were resuspended in PBS and serially diluted and plated onto selective LB media to differentiate and enumerate attacker and prey CFUs.

### CdiA-CT Toxin Domain Expression

*E. coli* T7 Express lysY/Iq cells bearing pESUMO expression vectors were grown shaking overnight in M9ZB + glucose media at 37°C. On the next day, overnight cultures were diluted 1:100 into M9ZB + glycerol media and grown shaking at 37°C until the OD600 reached 0.6-0.8. Expression was then induced with 250 μM IPTG and the temperature and shaking rate were reduced to 18°C and 180 rpm, respectively, overnight. Cells were harvested by centrifugation at 5000 x g for 10 minutes at 4°C, washed, and stored at -80°C until purification.

### Protein Purification

Frozen bacterial pellets were resuspended in 25 mL/L of culture of lysis/wash buffer (20 mM HEPES (pH 7.50), 500mM NaCl, 20mM imidazole, 1mM DTT, 1mM phenylmethylsulfonyl fluoride (PMSF)). Cells were lysed by sonication using a Branson Sonifier S-450 cell disruptor (Branson Ultrasonics Corp., Danbury, CT) at 70% amplitude for a total of 2 minutes with 30 second pulses, separated by 10-minute rest periods in an ice bath. Lysates were clarified by centrifugation at 13,000 x g for 45 minutes at 4°C. The supernatant was filtered using 0.45μm filters before being applied to Ni-nitrilotriacetic acid (NTA)-agarose beads (Bio-Rad, Hercules, CA) by gravity flow to isolate His_6_-tagged CT domains. Nonspecific proteins were removed with 100 mL lysis/wash buffer. Bound immunity factors were denatured from the CT domains with 8M guanidine-HCl, then eluted in 20 mL of lysis/wash buffer with 250mM imidazole.

Individual 1 mL elution fractions were pooled and purified proteins refolded by dialysis against 3L lysis/wash buffer without imidazole overnight at 4°C. The dialyzed protein was brought to room temperature before cleavage of the His_6_-SUMO tag by addition of 0.1 mg/mL Ulp1 protease for one hour at room temperature, then at 4°C overnight.

Cleavage reactions were reapplied to Ni-NTA resin by gravity flow to remove His6-SUMO/His6-Ulp1 (cleaved protein is collected in the flowthrough). The cleaved protein was then concentrated, assessed by SDS-PAGE and Coomassie staining for purity, flash frozen in liquid nitrogen, then frozen at -80°C until use.

### Nuclease Activity Assay

*In vitro* nuclease activity was assessed in 1X reaction buffer (20 mM HEPES (pH 7.50), 150 mM NaCl, 1 mM DTT, 2 mM MgCl_2_ or 2 mM CaCl_2_). Purified total 500 ng RNA or pUT18C plasmid DNA from *E. coli* strain DH5α was incubated with increasing amounts of CT domains of interest at 37°C for 30 minutes. Following incubation, DNA cleavage reactions were run on a 1% agarose TAE gel for 20 minutes at 120V before staining with SYBR Gold. Total RNA cleavage reactions were run on 1X TBE 10M urea 15% acrylamide gels for 40 min at 200V before staining with SYBR Gold. All gels were imaged using an Azure 400 imager (Azure Biosystems Inc., Dublin, CA).

### Deaminase Activity Assay

*In vitro* deaminase activity was assessed in 20 mM HEPES pH 7.50 150 mM NaCl 1 mM DTT. ssDNA primers with 6 FAM 5’ ends or dsDNA primers with unlabeled complementary strand were incubated with purified CT domain 3 for 1 hour at 37°C. Following incubation 5 μl of UDG reaction solution (Thermo Fisher, 0.02 U μl-1 UDG in 1X UDG buffer) was added to the deaminase reaction and incubated for 30 minutes at 37°C. To cleave abasic sites generated by UDG cleavage of uracil 100 mM NaOH was added and incubated at 95 C for 2 min with immediate addition to ice. Reactions were run on 1xTBE 10 M Urea 15 % acrylamide gels for 40 min at 200 V imaging on an Azure 400 imaging system (Azure Biosystems. Inc, Dublin, CA).

### Ethics Statement

All procedures were performed in accordance with the guidelines approved by the Loyola University Chicago Animal Care and Use Committee as described in protocol 24-022. Six to seven-week-old male and female BALB/c mice were purchased from The Jackson Laboratory (Bar Harbor, ME) and were housed under ABSL-2 containment at the Loyola University Chicago Center for Translational Research and Education Comparative Medicine Facility.

### Murine Pneumonia Bacterial Competition Assays

Overnight cultures of *P. aeruginosa* were diluted 1:50 into fresh LB supplemented with appropriate antibiotics and grown to mid-exponential phase. Strains were normalized to an infectious dose of 1x10^8^ CFU, pelleted, and resuspended to equal ratios in PBS. For infections using formalin-killed bacteria, 1x10^8^ CFU were fixed for 30 minutes in 10% formaldehyde at room temperature with gentle mixing. After fixation, cultures were washed three times with PBS, then resuspended to the appropriate volumes. Pulmonary aspiration was performed with 50 μL of bacterial suspension to promote inhalation of bacteria into the lungs. Mice were monitored for acute distress and disease during the five-hour experiment. After five hours, the mice were humanely euthanized and the lungs removed and homogenized for bacterial enumeration.

## Acknowledgements

We thank members of the Hullahalli Lab for their critical analysis of the experimental data and manuscript.

## Author Contributions

N.P., B.L.C., and J.P.A. conceived the study. N.P., B.L.C., and A.S.B. completed the experiments and analyzed data. N.P., A.S.B., and J.P.A. wrote the paper.

## Supporting Information

**S1 Figure.**
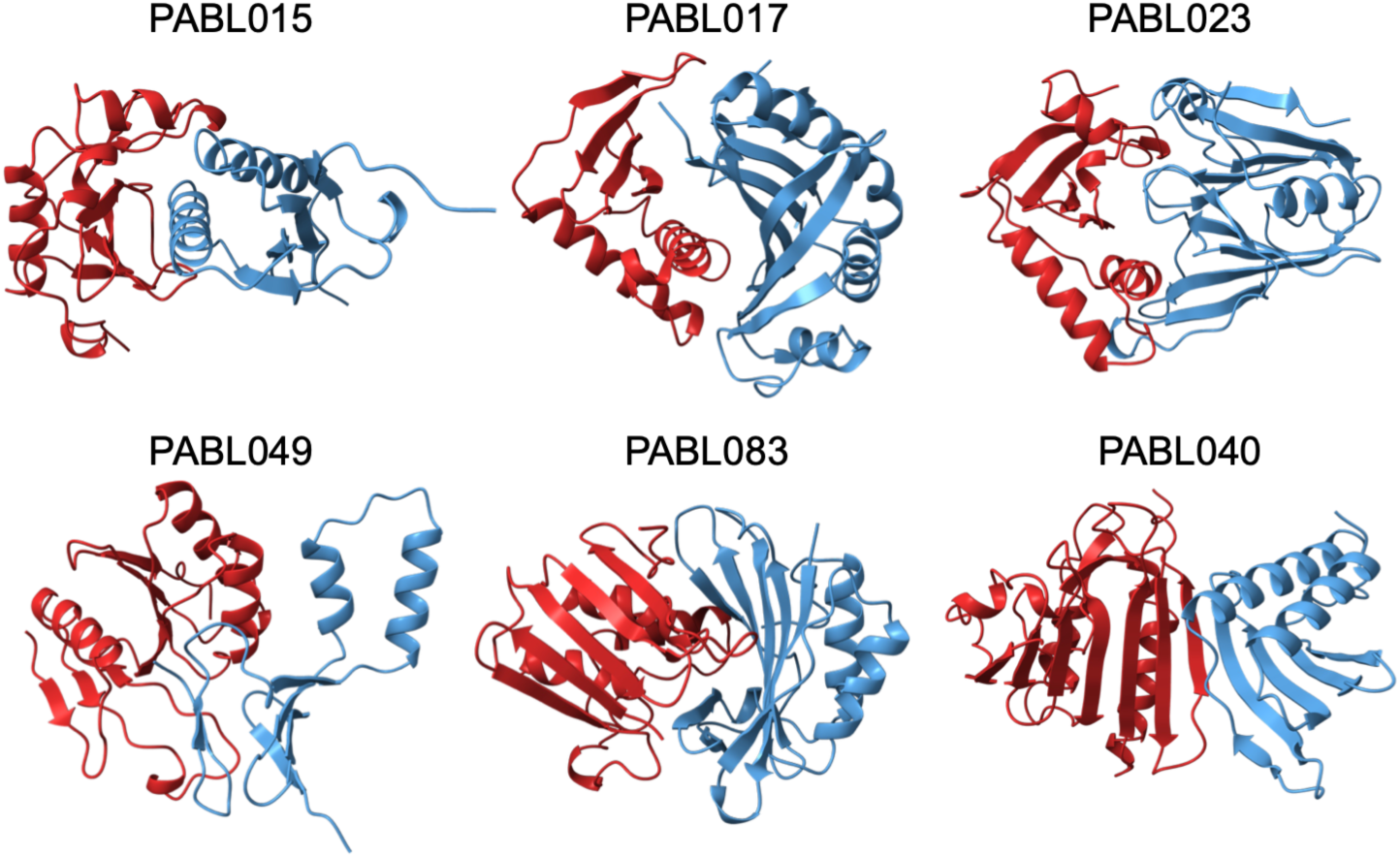
Predicted structures of CdiA toxin domains and CdiI Immunity proteins from this study. AlphaFold3 was used to predict the structure of the CdiA-tox and CdiI complexes for the different alleles used in this study. CdiA toxin domains are colored in red and associated CdiI immunity proteins are colored in blue. The representative strain containing each complex is noted above.

**S2 Figure.**
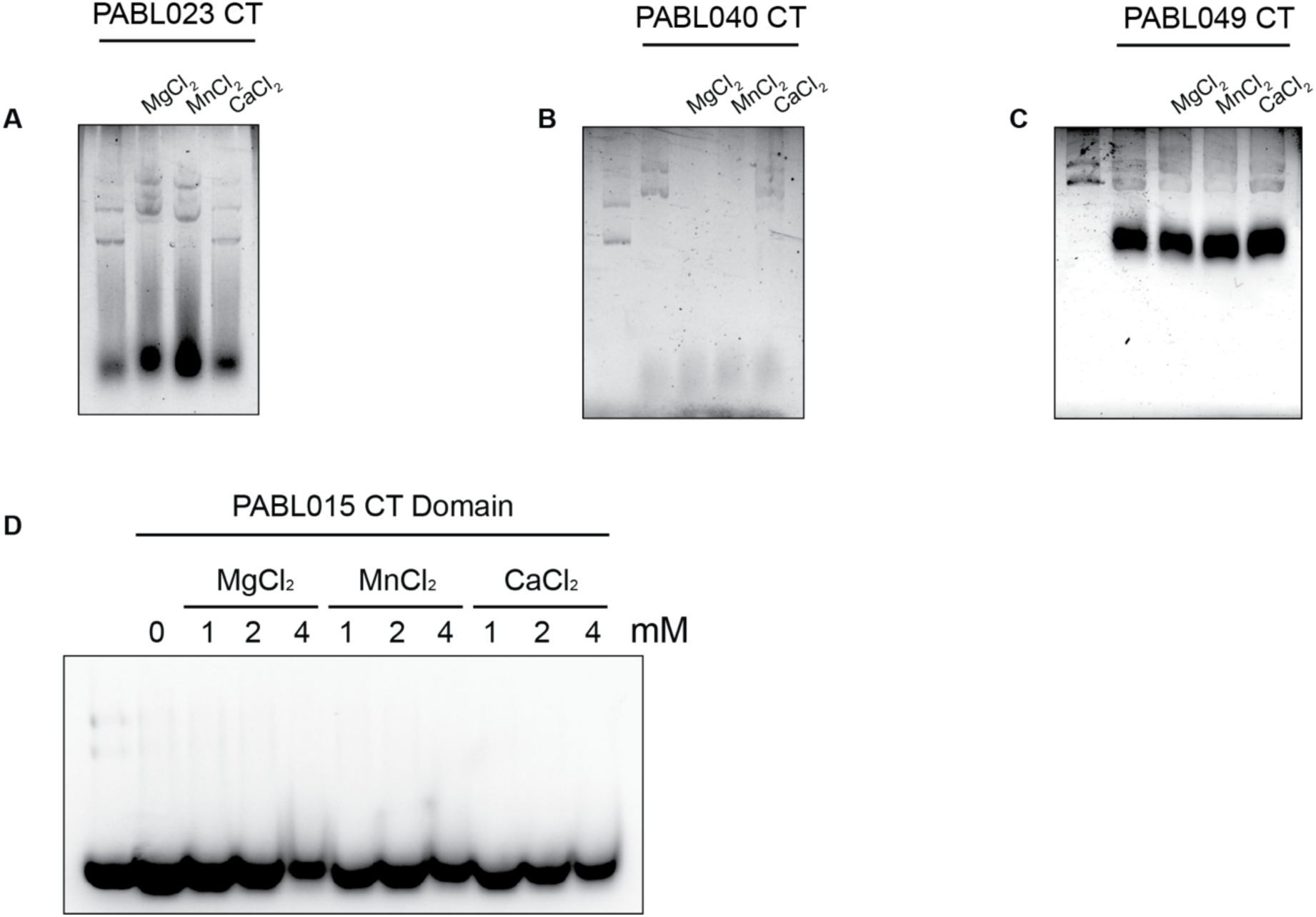
In vitro nuclease activity of CdiA toxin domains (CTs) from *P. aeruginosa* isolates. CdiA-CTs from strains PABL015, PABL023, PABL040, and PABL049 were purified independently (in the absence of CdiI immunity proteins) and assayed for nuclease activity. Reactions contained 10 µM toxin domain and either pUT18C plasmid DNA (A–C) or E. coli DH5α bulk RNA (D) as substrates. Activity was assessed by agarose gel electrophoresis following incubation in the presence of 2 mM MgCl₂ , MnCl₂ , or CaCl₂ . Substrate depletion and product cleavage were monitored relative to controls lacking divalent cations ("no metal") or toxin domain ("no toxin").

**S3 Figure.**
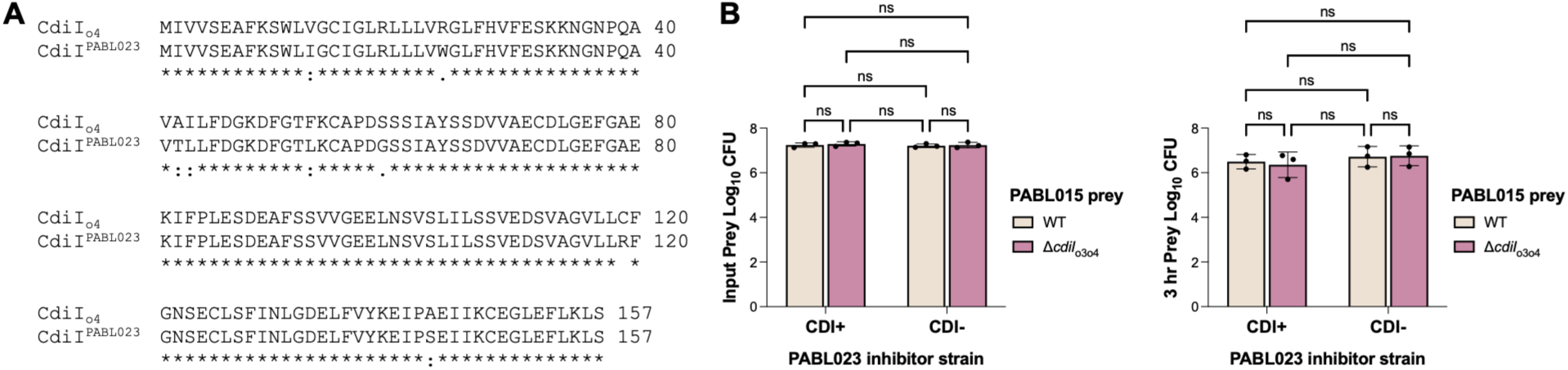
Lack of PABL023 CDI against PABL015 is not due to PABL015 CdiI_04_. Multiple sequence alignment of PABL015 CdiI_04_ with CdiI^PABL023^. Asterisks represent identical residues. (B) Mixed-strain competitions were performed using different PABL015 prey strains against CDI+ or CDI-PABL023 strains. The initial numbers of PABL015 prey bacteria at the beginning of competition (Input) and surviving prey after competition (3 hr) were quantified by plating serial dilutions on selective media. Data represent the mean ± SD from three independent experiments performed in triplicate. PABL015 WT and Δ*cdiI*_o3o4_ prey strains are colored tan and pink, respectively. Statistical significance between different prey strains was evaluated using a Two-Way ANOVA with Tukey’s multiple comparison test (ns, not significant).

**S1 Table. Strains used in this study.**

**S2 Table. Plasmids used in this study.**

**S3 Table. DNA oligo sequences.**

